# In situ GCIB Cryo-Sectioning Enables Subcellular Cryo-ToF-SIMS Imaging of *Arabidopsis* Seeds

**DOI:** 10.1101/2025.11.28.691143

**Authors:** Claire Seydoux, Bérangère Moreau, Michel Boujard, Eric Gautier, Pierre-Henri Jouneau, Jean-Paul Barnes

## Abstract

Time-of-flight secondary ion mass spectrometry (ToF-SIMS) enables label-free molecular imaging at submicrometric resolution, but its application to biological samples remains limited due to sample preparation challenges. Conventional fixation or dehydration alters morphology and induces analyte relocation, while cryo-transfer systems are costly and technically demanding. We present an *in situ* cryo-etching approach using a gas cluster ion beam (GCIB) and a new sample holder with a flat titanium ridge mask, enabling the sectioning of frozen specimens directly inside the ToF-SIMS instrument. Using *Arabidopsis thaliana* seeds as a model, this method produced flat, artefact-free surfaces suitable for subcellular imaging without chemical treatment or cryo-transfer. Mass spectra showed intact molecular profiles up to 1000 Da, and ToF-SIMS ion maps revealed preserved tissue architecture and distinct subcellular compartments at a resolution of ∼1 µm. Compared to air-dried cryosections, cryo-etching eliminated structural collapse and analyte delocalization. This workflow provides a practical and accessible route for cryo-ToF-SIMS analysis of hydrated biological materials, combining structural fidelity with molecular integrity. It offers a simple alternative to conventional cryo-transfer methods for high-resolution chemical imaging.

## INTRODUCTION

Over the past two decades, the development of new cluster ion sources (Au, Bi, Ar, and water GCIB) has enabled the analysis of complex molecules in biological and organic materials by time-of-flight secondary ion mass spectrometry (ToF-SIMS).^1^ Modern instruments can typically achieve lateral resolutions down to ∼50 nm using Bi cluster sources.^2^ However, such resolutions are rarely reported in current life science studies, which generally focus on tissue- or cell-scale imaging. A major reason for this is that high resolution requires careful beam alignment and reduced beam current, while acquisition times increase quadratically with reduced pixel size, making subcellular imaging particularly time-consuming.

Another crucial limitation hampering very fine resolution on biological specimens is sample preparation. Indeed, a prerequisite for accurate subcellular imaging is maintaining both the structural and chemical integrity of the specimen down to the desired resolution.^3–6^ Conventional protocols involving chemical fixation and/or dehydration by various means can disrupt cellular integrity,^7^ leading to cell deformation or analyte relocation artefacts.^8^ Analyte diffusion differs from molecule to molecule,^9^ making it difficult to verify that no such diffusion occurs in practice when studying a wide range of molecules.

When combined with appropriate sample preparation procedures, cryogenic methods, originally developed for cryo-electron microscopy, allow analysis of frozen-hydrated samples while preserving the physical structure of the specimen and preventing molecular migration or relocation. Although cryo-ToF-SIMS has become increasingly popular,^10–14^ its wider adoption remains limited by the need for dedicated cryotransfer systems, which are not widely available in most laboratories.

In this study, we propose a new sample preparation method for cryo-ToF-SIMS based on *in situ* cryo-etching using a gas cluster ion beam (GCIB) and a flat titanium ridge mask. This approach enables flat sectioning of the specimen directly inside the ToF-SIMS instrument. The sample, in the present case a 500 µm x 300 µm ovoid-shaped *Arabidopsis thaliana* seed, is then oriented perpendicular to the analyzer for optimal extraction and imaged by ToF-SIMS, demonstrating subcellular resolution. Results show superior structural and chemical preservation compared to conventional dehydrated preparations. This paper is based on a pending patent.^15^

## EXPERIMENTAL SECTION

### Cryo-microtomy and thaw mounting for reference imaging

Thaw-mounted sections were used to obtain reference ToF-SIMS images of dehydrated sections of *Arabidopsis thaliana* seed sections. Seeds were imbibed overnight in deionized water to soften their testa, then embedded in optimal cutting temperature (OCT) compound and frozen at -20°C. Sections of 5 µm and 20 µm thickness were cut at -20°C using a cryomicrotome (Leica CM1860). Sections were deposited on indium tin oxide-coated glass slides with a fine brush and allowed to air-dry at room temperature.

### Sample holder design for in situ cryo-etching

A custom titanium holder was fabricated to enable *in situ* GCIB sectioning. This holder features a small groove to secure seeds in front of a sharp titanium edge that serves as a mask for the GCIB. After GCIB etching, the holder can be rotated to expose the etched face to the ToF-SIMS analyzer.

For this realization, a 6 mm thick titanium plate was shaped in-house using electrical discharge machining (Figure S1). The rough surface near the groove was mechanically polished using diamond polishing discs down to 6 µm grit (Figure 1A). The tip of the holder was then cut with a wire saw to prevent collision with the analyzer once inserted into the ToF-SIMS instrument. Finally, the inner edge of the groove was polished using a xenon plasma focused ion beam (p-FIB, Helios 5, Thermo Fisher Scientific) at 30 kV, 500 nA, and 100 µs dwell time, over several boxes of 667 x 31 µm with 85% overlap (Figure 1B). A 2° angle was applied between the edge and the FIB beam to minimize redeposition. SEM imaging was used to verify the edge sharpness (Figure 1C).

**Figure 1.**
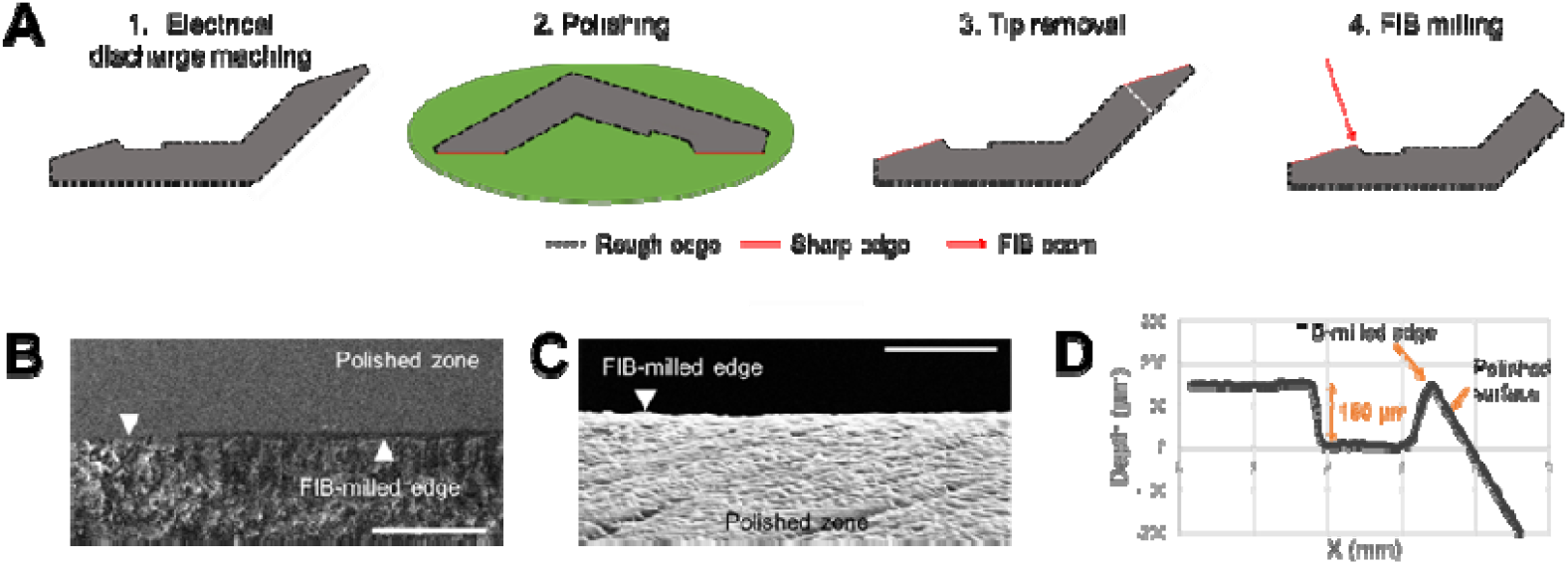
Fabrication and characterization of the sample holder. (A) Schematic of the manufacturing process. (B) SEM image of the edge during FIB milling, viewed from the top. Scale bar: 200 µm. (C) SEM image of the edge after FIB-milling, viewed from the side. Scale bar: 2 µm. (D) Topographic scan of the sample holder surface.

### In situ cryo-etching and imaging

Imbibed seeds were placed in the groove of the sample holder and frozen inside the ToF-SIMS loading chamber (NanoTOF II, ULVAC PHI) under nitrogen flow at atmospheric pressure. After cooling to -100 °C, the introduction chamber was pumped to 10^-5^ Pa and the sample was transferred into the main chamber. Seeds were etched by sputtering with a 20 kV Ar_2500_^+^ GCIB beam (60-80 nA) in non-interlaced mode, using 50 seconds sputtering cycles followed by 10 seconds charge compensation with a 1.5 kV oxygen gun and a 30 eV electron gun (Figure 2).

**Figure 2.**
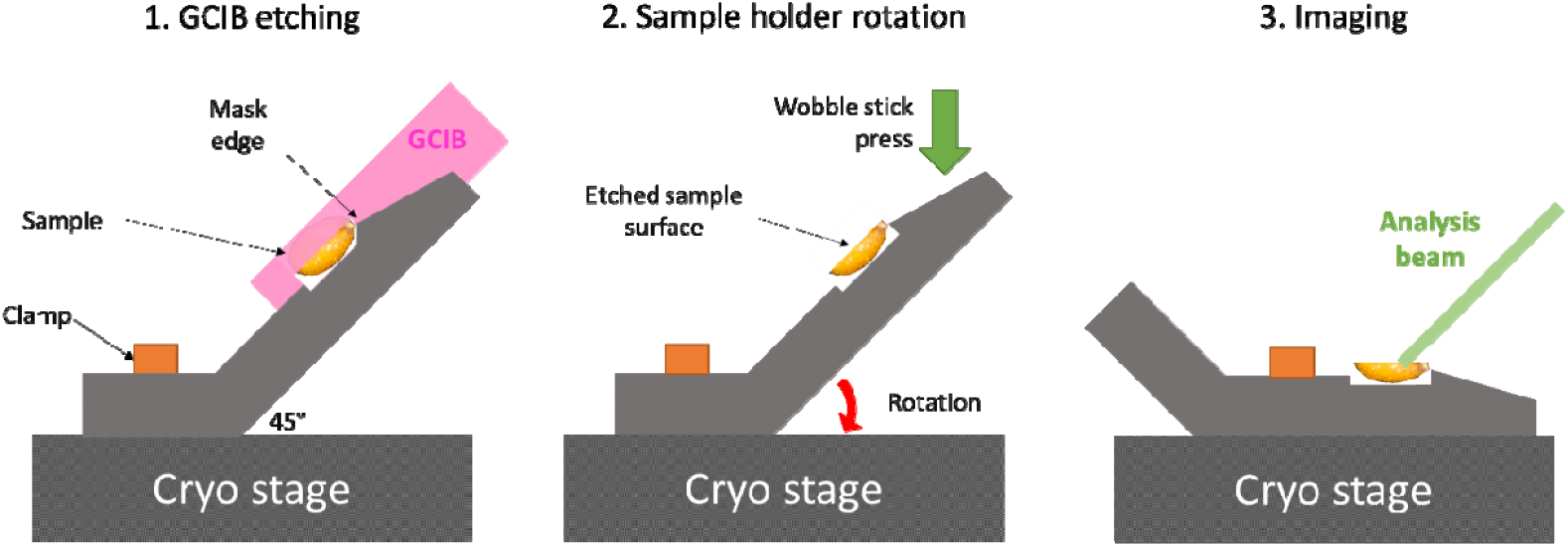
Operating principle of the specimen holder for cryo-abrasion and then cryo-imaging.

After etching, the holder was returned to the introduction chamber. A wobble stick was used to press the sample holder and rotate it into the analysis position (Figure 2). The sample was then reintroduced into the main chamber for imaging. Surface frost was removed by 10 minutes of gentle sputtering using the GCIB at 3 kV, 2 nA over a 800 x 800 µm raster area.

Seeds were imaged with a 30 keV Bi_3_^2+^ beam. Charge compensation was achieved using an electron flood gun (both polarities) and a 1.5 keV oxygen gun (positive polarity only). Between imaging frames, soft sputtering was applied using a 2.5 keV Ar_2500_^+^ GCIB in non-interlaced mode.

### Data analysis

Data were corrected for lateral drift using TOF-DR 3.1.0.10. Peaks were selected, and the ion images were exported from TOF-DR in .*tif* format (which preserves counts as long as they are below 255). Denoising was performed using PCA-assisted Noise2Void (PCA-n2v) as described by Seydoux et al.^16^

### Post-analysis characterization

After analysis, samples were freeze-dried in the instrument as it returned to room temperature. Samples were then imaged by scanning electron microscopy (Zeiss GeminiSEM 460) using the variable-pressure (low-vacuum) mode at 20 Pa, 5 kV, 1 nA, to assess the quality of the etched surfaces and structural damage. Topographic measurements of an etched seed were also performed using an optical microscope (ZEISS Axio Imager M2m) with the extended depth-of-field mode.

## RESULTS AND DISCUSSION

### Limitations of thaw-mounted and air-dried samples

Cryo-microtomy remains the most common sample preparation method for biological analysis by ToF-SIMS. Typically, samples are embedded in a medium, frozen, and sectioned into 2 to 50 µm slices using a sharp razor blade in a cryomicrotome. The sections are then mounted on a conductive slide and freeze-dried or simply air-dried.

Even if the morphology of seeds that were thaw-mounted and air-dried appeared acceptable, thicker sections of 10 µm and 20 µm exhibited strong ion suppression and topographical irregularities (Figures 3A and 3B, white arrows). Only 5 µm sections were sufficiently flat to avoid high topographic artefacts and ion suppression effects, permitting ion images of sufficient quality (Figure 3C). This is likely due to the very heterogeneous nature of plant tissues, particularly seeds, where individual cells are enclosed by cellulosic walls and the seed itself by a thick, rigid coat. The high topography observed could therefore result from collapse of the cellular contents during drying, while cell walls remain in place.

**Figure 3.**
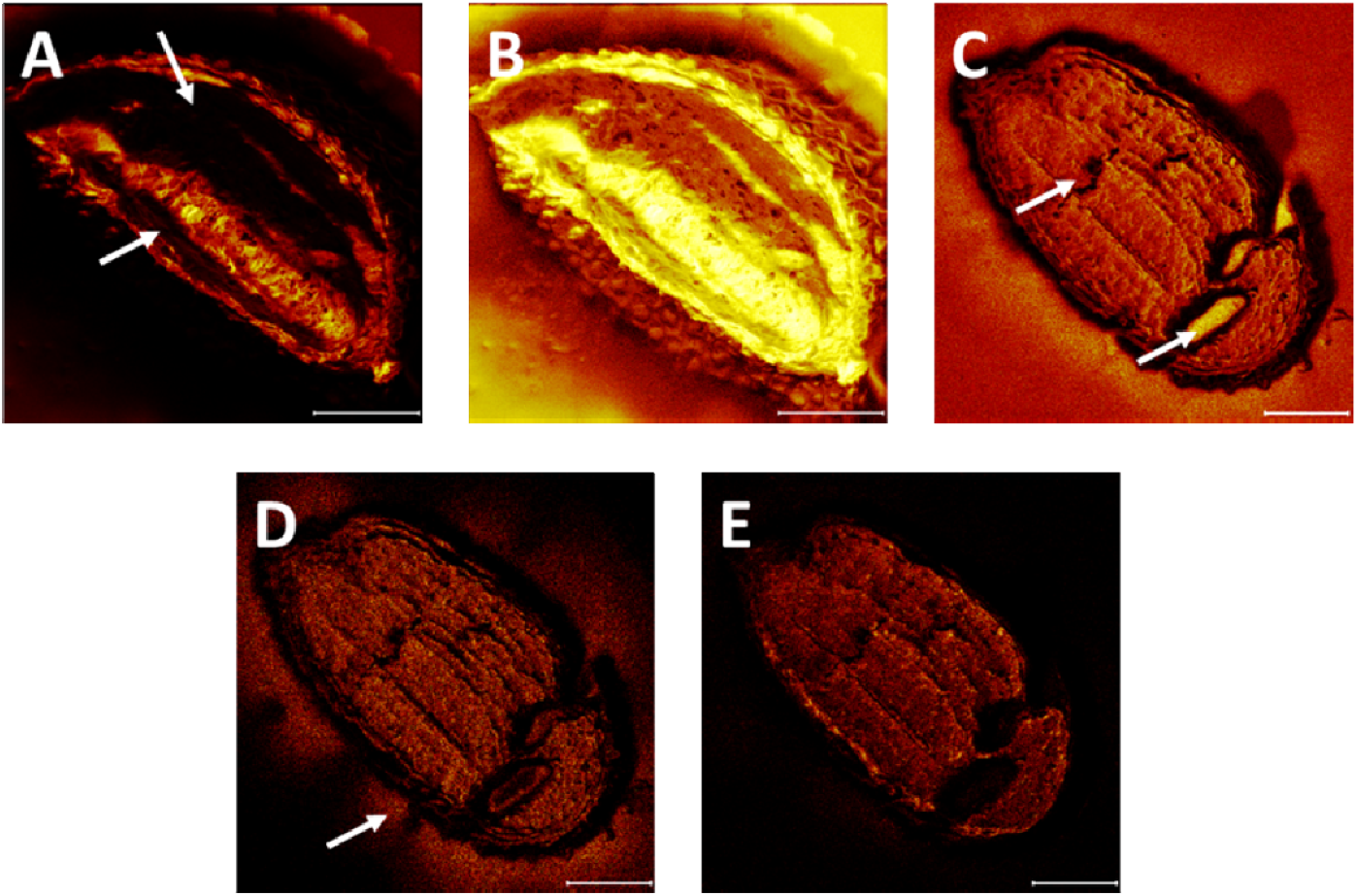
ToF-SIMS images of thaw-mounted and air-dried *Arabidopsis* seed sections. (A) Total ion image of a 20 µm-thick section. (B) Same image in logscale. (C) Total ion image of a 5 µm-thick section. (D) Ion image of C_18_H_31_O_2_^-^ at m/z = 279.23. (E) Ion image of PO_3_^-^ at m/z = 79.96. Scale bars: 100 µm.

The morphology of the 5 µm-thick seeds was overall satisfactory. However, significant structural damage was apparent (Figure 3C, arrows). ToF-SIMS analysis revealed substantial analyte delocalization: the fatty acid linoleate was detected both inside and outside the seed (Figure 3D), strongly suggesting fatty acid leakage. Interestingly, although all fatty acids were detected with the same pattern as linoleate, not all analytes leaked out of the seed. For instance, the phosphate ion PO_3_^-^ was detected exclusively inside the seed (Figure 3E). This may be explained by the fact that phosphate is a component of proteins and other insoluble contents, which are less prone to migration than triacylglycerides, which constitute most of the lipids in *Arabidopsis* seeds.^17^

In summary, due to structural damage and analyte delocalization, thaw mounting followed by air drying appears to be an unsuitable strategy for metabolic imaging of *Arabidopsis* seeds.

### Design and performance of the in situ cryo-etching holder

Cryo-microtomy methods hence appear inadequate for subcellular imaging of seeds, likely because of the difference in roughness between the tegument and the interior of the seed. Frozen-hydrated analysis might therefore be necessary for subcellular analysis of *Arabidopsis* seeds.

Previous studies have explored the possibility of performing *in situ* sample preparation using ion beams integrated into ToF-SIMS instruments. While Ga^+^ beams, now available on most instruments, allow for very precise sectioning,^18^ they have limited etching speed, making them unsuitable for objects with dimensions of hundreds of microns. Gas cluster ion beams, on the other hand, appear more attractive due to their high sputtering efficiency for organic materials.^19^ They are often used either as analytical beams or as a secondary beam to mitigate molecular damages from the primary imaging beam and perform depth profiling. Recent applications have demonstrated that these beams can also be used as etching beams for *in situ* preparations, while preserving the molecular integrity of the underlying layers.^20,21^ Besides, the sputter yield of argon GCIB for inorganic material is considerably slower than that of organic material.^19^

We took advantage of this property to design a custom titanium holder specifically meant for *in situ* etching of frozen *Arabidopsis* seeds. This sample holder features a titanium mask edge that enables flat etching by GCIB. This edge is 150 µm high, enabling a transverse cutting of the seeds at around the center of their large axis. To obtain a large field of view and optimize secondary ion extraction, the holder can be rotated by 45 degrees after cutting, as previously proposed,^18^ thus providing a flat surface over almost the entire seed, perpendicular to the analyzer extraction field.

This setup allows for the etching of one seed within a few hours (Figure 4A). Curtaining artefacts parallel to the beam are visible both in the electron microscopy images of the seed and in the total ion image (Figures 4B, 4C), likely due to imperfections in the polished edge. However, these did not result in major topography-related artefacts, unlike those previously observed (Figure 3). Furthermore, topological measurements (Figure S2) suggest that height differences on the etched surface of a seed remain below 5 µm. Most likely these imperfections are below the micrometer scale, as the rugosity of the edge roughness never exceeded this value. This conclusion is supported by the absence of topography-related artefacts on the ion image of the part of the seed that was already fully-etched.

**Figure 4.**
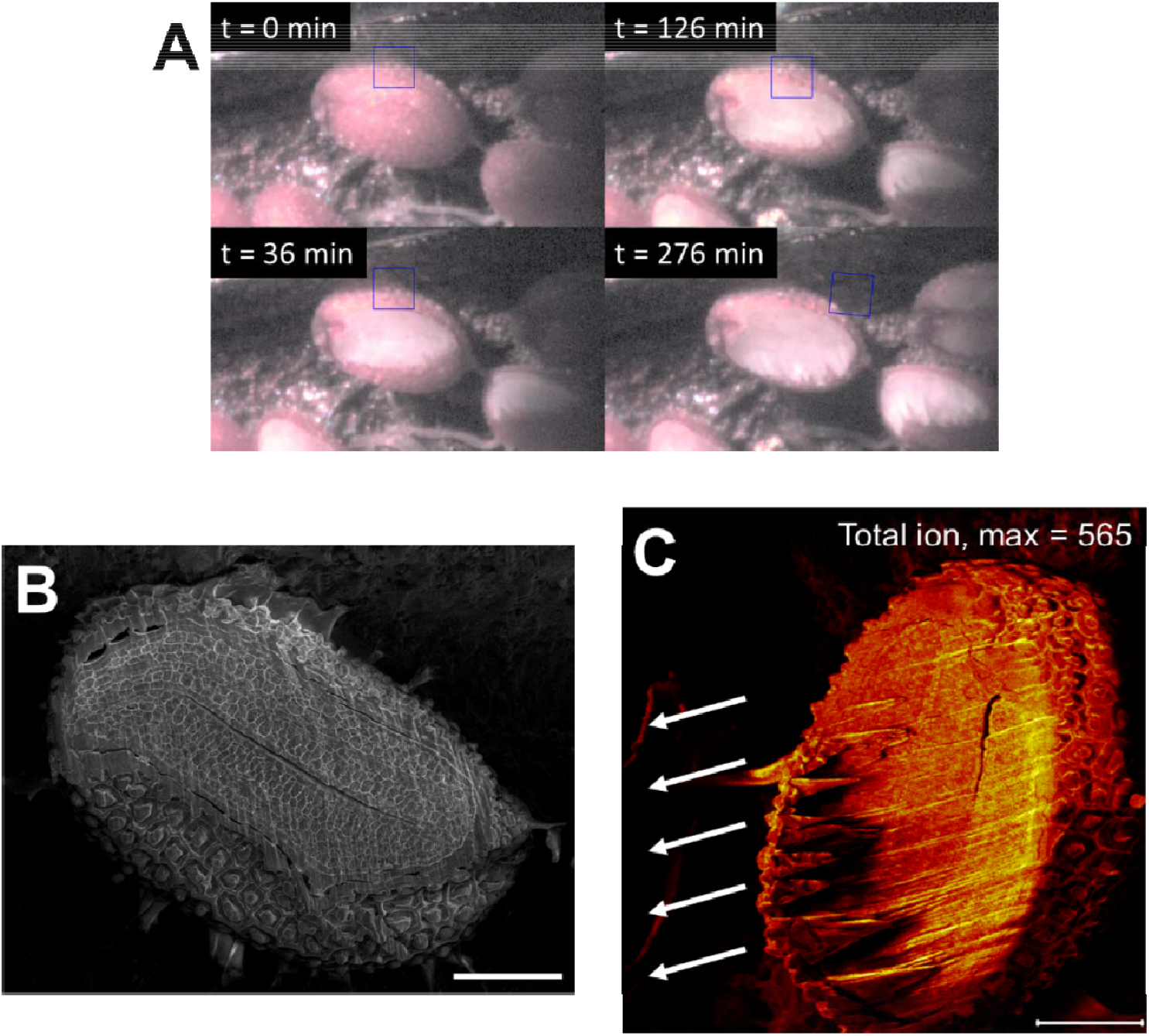
In situ sectioning of *Arabidopsis* seeds. (A) Optical images during the GCIB etching. Blue squares are 100 µm wide. (B) SEM image of an etched seed. (C) Total ion image of the same seed by ToF-SIMS. Arrows indicate the GCIB direction during etching.

### Subcellular molecular imaging in seeds after in situ cryo-etching

Having demonstrated that *in situ* cryo-etching of seeds produces sufficiently flat surfaces for ToF-SIMS imaging, two questions remain regarding the applicability of this method for subcellular imaging: *(i)* does high-energy (> 60 keV) GCIB induce molecular damage, and *(ii)* to what extent the localization of molecules is preserved?

The mass spectrum collected from an etched seed (Figure 5A) revealed four main regions: *(i)* in the m/z range below 150 Da, the usual low-mass fragment; *(ii)* in the range of 200-300 Da, monoacylglycerols and fatty acids; *(iii)* at 500-650 Da, diacylglycerols; and *(iv)* at 850-1000 Da, triacylglycerols. These closely match the lipid profiles previously identified from homogenized lipid extracts of *Arabidopsis* seed.^22^ This mass spectrum is also very similar to that of a seed prepared by cryo-microtomy (Figure 5B). These results suggest that *in situ* cryo-etching using a high-energy and high-intensity GCIB does not cause significant molecular degradation, at least below 1000 Da, and is thus fully compatible with ToF-SIMS molecular imaging.

**Figure 5.**
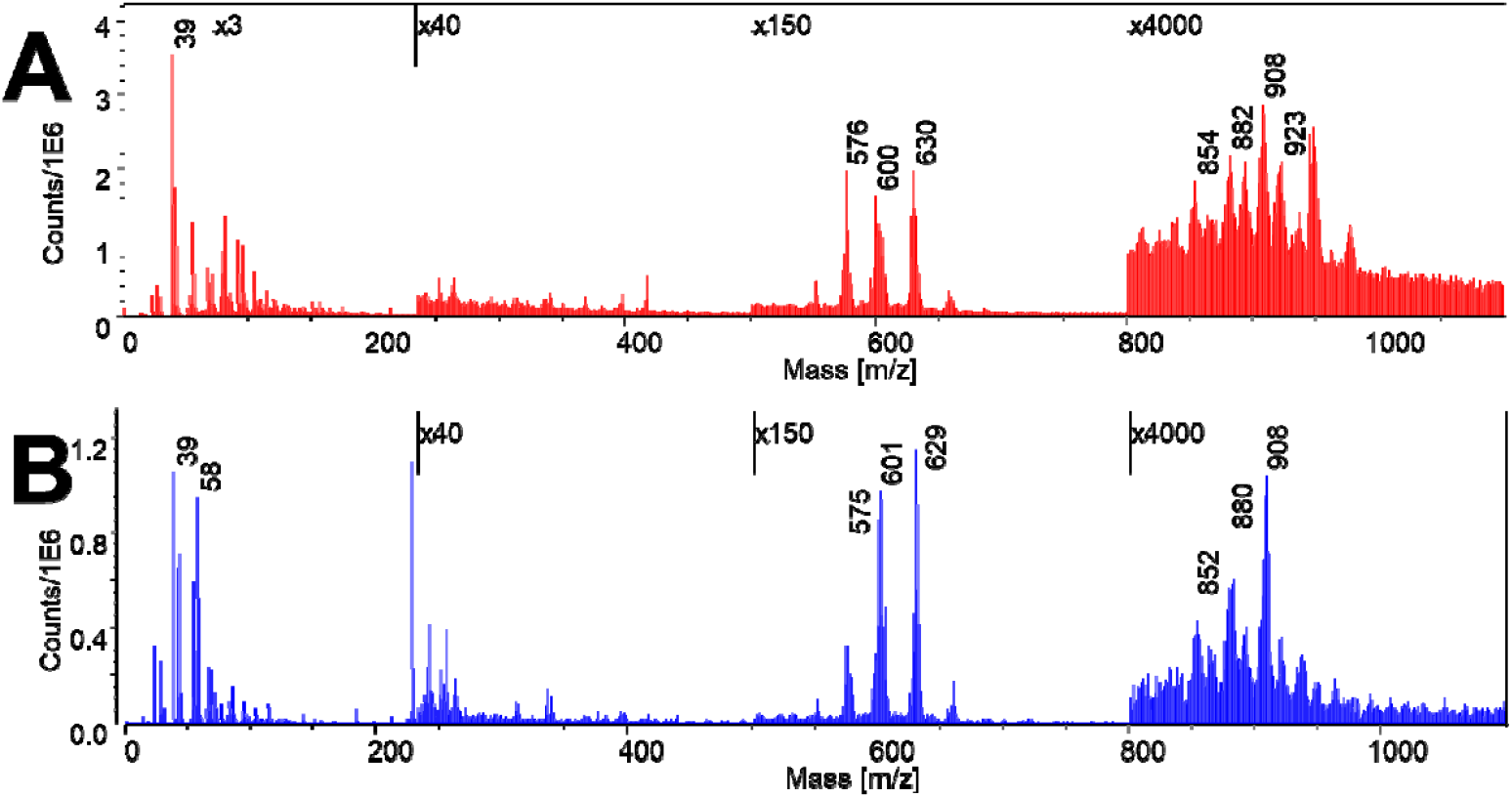
Mass spectra of *Arabidopsis* seeds obtained after (A) in situ cryo-etching and (B) cryo-microtomy. Expansion factors are indicated above.

Another requirement for this method to be suitable is that it should preserve the structure of the seeds and the localization of the molecules. Overlay images of sodium (m/z = 23, Na+), a protein fragment (m/z = 58, C3H8N+), and choline (m/z = 104, C5H14NO+) revealed a tissue architecture very similar to the one observed by microscopy in cross-sections (Figure 6B). The albumen, consisting of a single-cell layer in *Arabidopsis thaliana* seeds, is clearly visible, and individual cells are even identifiable, especially after PCA-n2v denoising (Figure S3). The albumen appears richer in choline than the embryo, which in turn is richer in protein bodies, consistent with their respective roles as regulator of seed germination and nutrient storage locus. While cryo-sectioning and air-drying did not allow us to distinguish the albumen from other tissues (Figure 3), in situ cryo-etching clearly preserves the separate identities of tissues at single-cell resolution, without leakage or homogenization.

**Figure 6.**
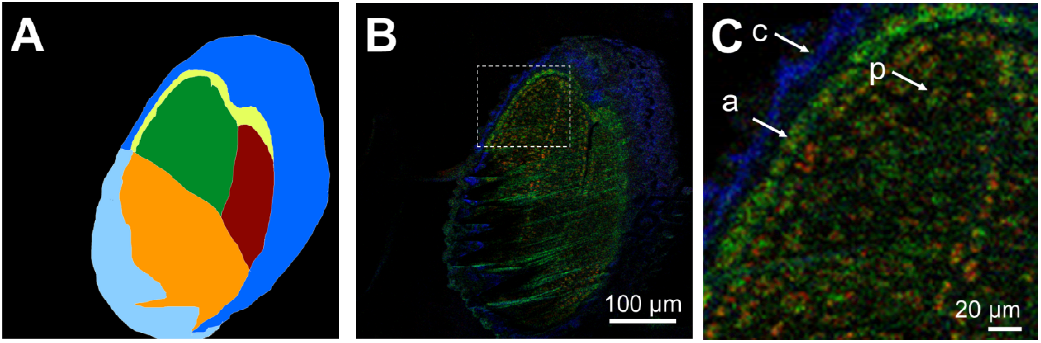
Subcellular mass spectrometry imaging of an *Arabidopsis* seed. (A) Image segmentation for interpretation: dark blue: seed coat, light blue: partially etched seed coat, yellow: albumen, green: cotyledons, red: radicle, orange: partially etched seed. (B) Overlay image of red: protein fragment, m/z = 58, green: choline, m/z = 104, blue: Na. (C) Inset from B. c: seed coat, a: albumen p: cotyledon protein body.

The lateral resolution obtained is approximately 1 µm, corresponding to the pixel size. However, for biological samples, the useful resolution is the minimum between the resolution effectively reached by the measuring beam and the scale at which the sample of interest is preserved. In our case, it appears that the subcellular architecture remained intact at the micron-scale. Cryogenic conditions prevent both chemical reactions and molecule movement and are accordingly a suitable solution for high resolution studies.

### Comparison of GCIB in situ cryo-etching with other methods

Compared to other preparation workflows, *in situ* GCIB cryo-etching offers several advantages and trade-offs.

Firstly, cryogenic GCIB etching preserves both molecular localization and chemical composition without embedding or dehydration where molecules are degraded. No molecular degradation or analyte relocation was observed, enabling subcellular imaging. While curtaining artefacts can appear, they did not significantly affect imaging quality. Furthermore, the gentle GCIB etching did not induce noticeable molecular damage in the mass range below 1,000 Da.

This workflow can be implemented on many currently available instruments, bypassing the need for an installation of a dedicated cryo-transfer system. High-pressure freezing and cryo-transfer systems remain the gold standard for ultrastructural preservation, especially for highly hydrated samples. This also also allows vitrification during freezing, which is not the case for slow freezing inside the loading chamber under atmospheric pressure nitrogen flow. However, these setups are costly, complex, and not accessible to many laboratories. The proposed method provides a practical alternative achieving adequate preservation for micron-scale imaging directly within standard ToF-SIMS instruments.

One major drawback is that *in situ* GCIB etching is time-consuming (typically several hours per sample) and requires operator skill. It can however conveniently be carried out overnight. Furthermore, frost formation and temperature control remain challenging. Each experiment requires careful handling to avoid contamination, but reproducibility is good once parameters are optimized.

Overall, *in situ* GCIB cryo-etching offers a practical, accessible approach for subcellular chemical imaging of complex biological specimens without the need for extensive cryo-infrastructure.

## CONCLUSIONS

Seeds are challenging samples to observe using high-resolution mass spectrometry imaging: their rigid outer layer makes sectioning difficult. To solve this problem, we developed a novel *in situ* preparation methodology involving freezing the sample within the instrument loading chamber, GCIB cross sectioning using a flat titanium edge as a mask, and subsequent reorientation for perpendicular imaging. This approach enables metabolic imaging while keeping the sample near its native state. Importantly, no physical or chemical degradation of the sample was observed.

Although demonstrated on seeds, the method can be adapted to a wide range of biological or organic specimens, if the sputtering yield of titanium remains negligible relative to that of the sample. The holder was designed to produce transverse sections of seeds, but its height can be easily modified to fit other types of samples. It would also be possible to combine the holder with a motorized rotation system, as previously proposed.^18^

In summary, this is a robust, and versatile workflow for high-resolution ToF-SIMS imaging of organic and biological materials.

## ASSOCIATED CONTENT

### Supporting Information

Figure S1, showing the exact shape used for electrical discharge machining of the sample holder, and Figure S2 showing the optical image and topological map of a seed after cryo-etching.

## AUTHOR INFORMATION

### Author Contributions

C.S., J-P.B., and P.-H.J. designed the experiments. C.S., B.M., M.B., and E.G. designed and fabricated the sample holder. C.S. and J.-P.B. and P.-H. J performed the ToF-SIMS and microscopy experiments. C.S., J.-P.B., and P.-H.J. wrote the manuscript. All authors approved the final version of the manuscript.

### Notes

The authors declare no conflict of interest.

## ACKNOWLEDGMENTS

CEA-Leti is a member of the Carnot institute network. This work was supported by the French National Research Agency (ANR) “3D-Lipid” InterCarnot project between CEA-Leti and 3BCAR institute. It was carried out on the Platform for Nanocharacterisation (PFNC) of CEA-Grenoble, supported by the “Recherche Technologique de Base” and “France 2030 - ANR-22-PEEL-0014” programs of the ANR.

---

**Figure S1.**
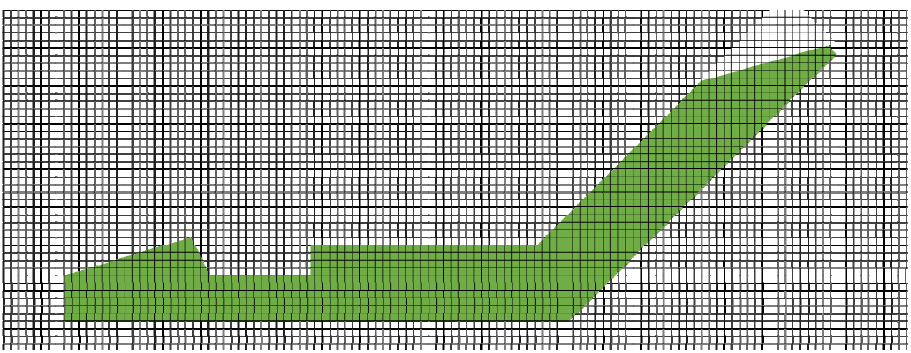
Side view of sample holder design for electrical discharge machining. Grid squares represent 100 µm. The thickness of the titanium plate was 6 mm.

**Figure S2.**
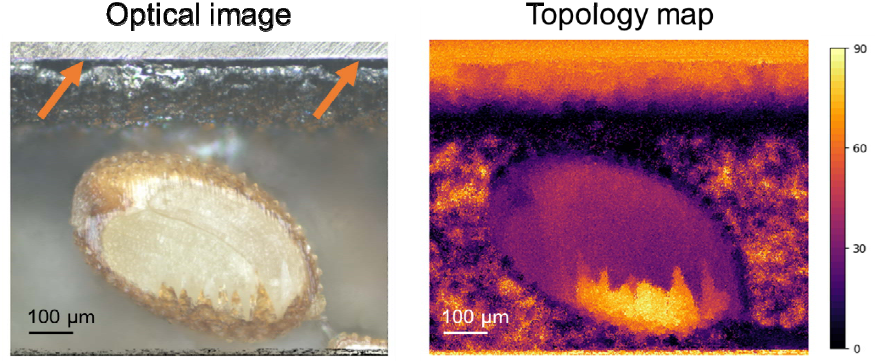
Optical imaging of seeds after cryo-etching. Orange arrows indicate the polishing edge. The background underneath the seed is out of focus, and the corresponding topology measurements are inaccurate and should be ignored.

